# Posterior parietal cortex activity during visually cued gait: a preliminary study

**DOI:** 10.1101/2025.04.10.648269

**Authors:** Paul McDonnell, Adam B. Grimmitt, Jonaz Moreno Jaramillo, Wouter Hoogkamer, Douglas N. Martini

## Abstract

Safe gait requires visually cued (VC) step adjustments for negotiating targets and obstacles. Effective step adjustments rely on good visuospatial processing. The posterior parietal cortex (PPC) is implicated in visuospatial processing, yet empirical evidence is limited for the PPC’s role during gait in humans. Increased cortical control of gait is associated with higher gait variability, a marker of gait performance and fall risk among older adults. However, the cortical underpinnings of gait variability in visually complex environments are not well established. The primary aim of this preliminary study was to assess PPC activity during VC gait and VC gait with perturbations (VCP). A secondary aim was to determine how PPC activity relates to gait variability during VC and VCP gait. Twenty-one healthy young adults completed three treadmill gait conditions at preferred speed: non-cued (NC) gait, VC gait, where stepping targets were presented in a regular pattern, and VCP gait, where stepping target positions were pseudorandomly shifted. Functional near-infrared spectroscopy quantified relative changes in deoxygenated and oxygenated hemoglobin (ΔHbO_2_) concentrations in the PPC. Inertial measurement units quantified gait variability. Moderate effects were observed for more positive ΔHbO_2_ from NC to both VC and VCP gait, likely reflecting the increased visuospatial processing demands. Stride time variability was positively correlated with PPC ΔHbO_2_ during VC gait, suggesting a potential role for the PPC in modulating temporal components of VC gait. Extending these findings to older adults will help to elucidate the PPC’s role in gait adaptability and fall risk with aging.

## Introduction

Everyday gait requires visually cued (VC) step adjustments, which are coordinated by cortical and subcortical pathways, to navigate complex gait environments. Mobile functional near-infrared spectroscopy (fNIRS) studies report increased prefrontal cortex (PFC) activity during tasks involving precision stepping (Koenraadt et al., 2014; Le et al., 2023) and obstacle negotiation (Chen et al., 2017; Mirelman et al., 2017, Pelicioni et al., 2022). Greater recruitment of PFC resources for VC gait contrasts with the limited PFC recruitment observed for non-cued gait, highlighting the complexity of the task. Though task performance may decline as task demands increase, this adaptive neural response reflects an effort to maintain performance (Reuter-Lorenz & Cappell, 2008). Unfortunately, the involvement of cortical regions beyond the PFC during VC gait remains comparatively understudied in humans. Emerging evidence indicates that the posterior parietal cortex (PPC) may play a critical role during VC gait (Drew et al., 2023; Nordin et al., 2019).

Interspecies studies implicate the PPC in planning VC gait modifications (Drew & Marigold, 2015; Nordin et al., 2019). The PPC processes and relays visual information concerning environmental targets or obstacles to guide the goal-directed trajectory and foot placement of the stepping limb (Buneo & Andersen, 2006; Marigold & Drew, 2017). Indeed, specific cells residing in the cat PPC encode either the time or distance to contact with an obstacle (Marigold & Drew, 2017). Using electroencephalography (EEG) in humans, Nordin et al. (2019) demonstrated that the PPC is reliably activated preceding the step over an obstacle during gait. During precision stepping paradigms with targets presented at fixed or variable distances, Yokoyama et al. (2021) reported higher PPC activity in the variable condition using EEG, while Le et al. (2023) reported no change in activity in the superior parietal lobule (a sub-region of the PPC) using fNIRS. A notable difference between the paradigms used by these authors was that the former manipulated both step length and width during the variable condition, while the latter manipulated step length only. Considering that mediolateral step adjustments represent a greater deviation from the habitual gait pattern (Hoogkamer et al., 2015), it is conceivable that these adjustments carry greater visuospatial processing demands, eliciting higher PPC activity. This idea aligns with suggestions that compared to anteroposterior gait adjustments, mediolateral gait adjustments are under more active control via sensory integrative feedback because the passive dynamics of gait are less stable in this direction (O’Connor & Kuo, 2009). Accordingly, the removal of visual information during gait disproportionately affects mediolateral gait dynamics compared to anteroposterior gait dynamics (Wuehr et al., 2013).

Gait variability (i.e., stride-to-stride fluctuations of spatiotemporal gait cycle parameters) is increasingly used as a marker for gait performance and fall risk (Lord et al., 2011). Gait variability is linked to gait automaticity (i.e., limited cortical control of gait), such that higher variability purportedly reflects reduced gait automaticity (Nóbrega-Sousa et al., 2020). Gait variability can differentiate older adults with and without mobility and cognitive impairment (Hausdorff, 1998; Verghese, 2008). Further, more variable stride time and stride length can predict falls in older adults (Hausdorff et al., 2001; Maki, 1997). That gait must dynamically adapt to negotiate targets and obstacles complicates the interpretation of gait variability in visually complex environments. Higher stride-to-stride variability can reflect this adaptability in response to environmental demands (Beauchet et al., 2009). Among both young and older adults, gait variability increases during obstacle negotiation (Mirelman et al., 2017; Nóbrega-Sousa et al., 2020). Moreover, positive associations between gait variability and fNIRS-measured PFC activity during obstacle negotiation (Mirelman et al., 2017) and during cognitive dual-task gait (Nóbrega-Sousa et al., 2020) highlight the importance of higher order brain regions for complex gait control. Unfortunately, beyond the above-mentioned associations with PFC activity (Mirelman et al., 2017; Nóbrega-Sousa et al., 2020), the cortical underpinnings of gait variability remain insufficiently characterized. Delineating the neural correlates of gait variability during tasks that mimic the visual processing demands of real-world gait could provide critical insights for neuromotor-based interventions aimed at reducing mobility impairment and fall risk in older adults.

While previous studies assessed PPC activity during VC gait, studies relating PPC activity to gait performance are scarce. Pizzamiglio et al. (2018) reported that during dual-task gait, increased PPC activity in young adults predicted lower mediolateral center of mass motion, suggesting a role for the PPC in regulating gait dynamics in complex gait environments. Moreover, considering that the PPC engages to register and store the spatiotemporal relationship between the body and environment (Marigold & Drew, 2017), there are likely important functional associations between PPC activity and gait variability. Indeed, magnetic resonance imaging studies report that reduced PPC grey matter volume (Beauchet et al., 2014) and integrity (Tian et al., 2016) are associated with increased temporal gait variability. Extending this work by using fNIRS to relate real-time PPC activity to gait variability during a VC gait task is a clear next step toward understanding the cortical mechanisms responsible for gait impairment and fall risk in clinical populations. Examining these relationships in young adults can provide useful baseline data to help better understand cortical deficiencies that occur across the lifespan.

The primary aim of this preliminary study is to determine PPC activity patterns during VC gait in young adults using fNIRS. A secondary aim is to determine how PPC activity relates to gait variability during VC gait. Since the PPC appears important for fast step adjustments (Potocanac & Duysens, 2017), and inaccurate reactive step adjustments relate to fall risk (Robinovitch et al., 2013), we use a VC gait task which requires reactive step adjustments. Our gait conditions comprise: 1) non-cued (NC) treadmill gait, 2) VC treadmill gait, where stepping targets follow a regular pattern, and 3) VC treadmill gait with perturbations (VCP), where stepping targets undergo pseudorandom position shifts. As with obstacle negotiation, cueing participants to make reactive step adjustments imposes gait variability. We therefore anticipate that gait variability measures will be highest during the VCP gait. We hypothesize that compared to NC gait, PPC activity will increase during VC gait and will increase further during VCP gait. Additionally, we hypothesize that increased PPC activity will relate to increased gait variability during both VC and VCP gait.

## Methods

### Participants

Twenty-one healthy young adults (22.2 [3.6] years, 8F) were enrolled after meeting the inclusion and exclusion criteria. Inclusion criteria comprised being 18-29 years old. Exclusion criteria comprised having a chronic musculoskeletal or neurological injury or disease; having a cardiovascular disease or arrhythmia; surgery that would affect gait within the past year; uncorrected vision impairment; or inability to understand written and spoken English. All participants provided informed consent before participating. A post-hoc power analysis was conducted using G*Power 3.1 to determine the achieved power for a repeated measures ANOVA, based on our observed medium effect size (*f*=0.35), alpha of 0.05, and a sample size of 21. Results indicated an achieved power of 0.93. This study was conducted in accordance with the Declaration of Helsinki and was approved by the Institutional Review Board at the University of Massachusetts Amherst (IRB #3501).

### Gait assessment

To quantify gait variability, participants were fitted with six body-worn inertial measurement units (IMUs) (Opal v2, APDM, a Clario Company, Portland, OR), sampling at 128Hz. IMUs were affixed to the feet, wrists, sternum, and lumbar spine (near the natural waistline, where a belt would be worn) using Velcro straps. IMUs were wirelessly synchronized through Mobility Lab ™ v2 software (APDM), which was used to initialize and collect the data. Gait measures reflect the full 3-minute walking period for each condition. The following outcomes measures were calculated by Mobility Lab™ proprietary algorithms: gait speed (m/s), stride length (m), stride length variability (m), stride time (s), stride time variability (s), lumbar mediolateral range of motion (RoM) (°), and lumbar mediolateral RoM variability (°). Variability measures were computed by Mobility Lab™ as the standard deviation of each measure across each walking condition, from which we calculated the coefficient of variation (CoV; (SD/mean) × 100) (%). CoV measures comprised the primary outcome measures from the gait assessment.

### Gait conditions

Participants completed three 3-minute gait conditions on a split-belt treadmill (Bertec, Columbus, OH) at self-selected speed in the order listed: NC gait, VC gait, and VCP gait. During the second and third condition, illuminated rectangular visual stepping targets, attuned to each participant’s foot size and average step length and width, were projected onto the treadmill belts, and approached the participant at belt speed (Figure 1). Participants were instructed to step as accurately as possible onto the targets, aiming to hit the center of each rectangular target with the imagined center of their foot sole. During the VCP condition, target shifts in the anterior, posterior, or lateral direction were pseudorandomly imposed every 3-7 steps, requiring a step adjustment. Target shifts occurred when the target came within 130% of the participant’s average step length (measured from a reflective motion capture marker worn on the sacrum) (Mazaheri et al., 2015). Targets were shifted by 40% of the participant’s average step length in the anterior or posterior direction, or by 20% of step length in the lateral direction (Mazaheri et al., 2015). Immediately prior to each 3-minute condition, there was a 20-second quiet standing period, serving as a baseline for cortical activity measures.

**Fig 1:**
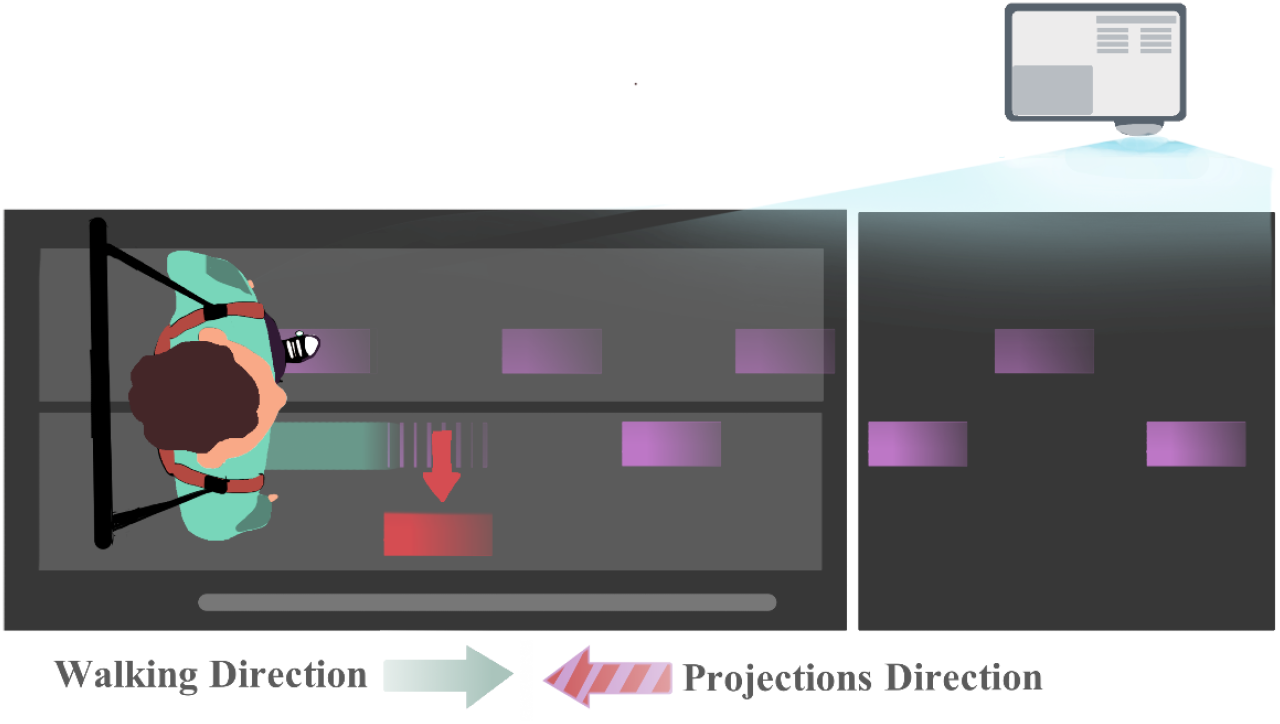
Schematic displaying the experimental set-up for the visually cued gait conditions. Participants walked on a split-belt treadmill while wearing a safety harness. Purple rectangles are stepping targets, attuned to each participant’s foot size and average step length and width, presented consistently during visually cued gait. The red rectangle is an example of a perturbed target during perturbed visually cued gait, undergoing a lateral positional shift, upon coming within 130% of step length (green) from a sacrum-worn reflective marker (not shown). Target positional shifts in the anterior-posterior direction were also imposed (not shown)

### fNIRS data acquisition

Relative changes in oxygenated hemoglobin (HbO_2_) and deoxygenated hemoglobin (HHb) were quantified during each gait condition with a portable, 24-channel continuous-wave fNIRS device sampling at 50Hz (Dual Brite MKII; Artinis Medical Systems, Netherlands). Twenty-two long channels (source-detector distance: 30mm) and two short-separation channels (source-detector distance: 10mm) were split evenly over the left and right hemispheres. After fitting the fNIRS cap according to each participant’s head circumference, a digitizer (Patriot 3D Digitizer, Polhemus, Vermont) was used to digitize optode positions based on anatomical references (nasion, inion, bilateral preauricular points, and vertex). Parietal positions P3 and P4 from the International 10-20 System were determined. P3 and P4 demonstrably correspond to Brodmann area 7 (Homan et al., 1987). Visual inspection of channel locations was performed in the Patriot system GUI. From the available channels, four (two per hemisphere) channels were selected for analysis, corresponding to the P3 and P4 positions (i.e., left and right PPC; Figure 2). Selected channels were consistent across all participants. Two short-separation channels were employed (one per hemisphere) positioned at T4 and T5 on the 10-20 system.

**Fig 2:**
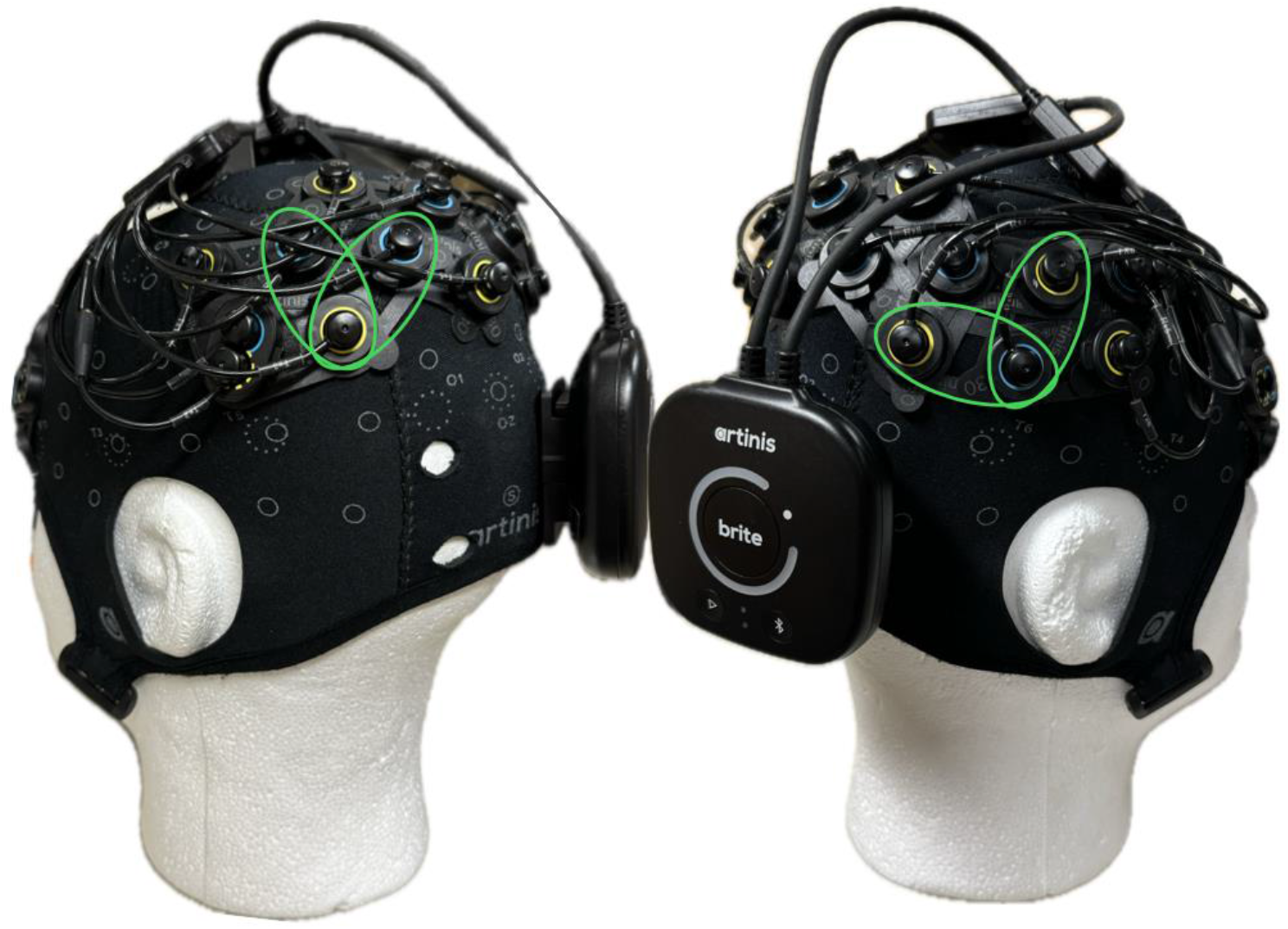
fNIRS optode configuration comprising 10 transmitters (yellow) and 8 receivers (blue). Key channels (transmitter-receiver combinations) selected for analysis are indicated with green ellipses

fNIRS data were processed in MATLAB 2023b. First, a processing pipeline involving selected functions from the HOMER3 package (Huppert et al., 2009) was applied. Raw data were visually inspected, and a channel pruning function (*hmr_PruneChannels*) was employed to flag channels with a signal to noise ratio of less than 5 (von Lühmann et al., 2020). Raw light intensity was converted to optical density (*hmr_Intensity2OD*). A motion artifact detection function (*hmr_MotionArtifactByChannel; tMotion = 1*.*0, tMask = 1*.*0, STDEVthresh = 40*.*0, AMPthresh = 5*.*0*) was then used to identify spike-like motion artifacts based on where the signal exceeded a threshold in change of standard deviation within a predefined time window (Huppert et al., 2009). This threshold was set at 40 after visual inspection of the dataset, to optimize artifact detection, following the same approach used by others (Cooper et al., 2012; Di Lorenzo et al., 2019). A cubic spline interpolation (*hmr_MotionCorrectSpline; p = 0*.*99*) corrected the artifacts flagged by the detection algorithm (Scholkmann et al., 2010). A wavelet filter was next applied (*hmr_MotionCorrectWavelet; iqr = 1*.*50*), in line with reports that combined spline and wavelet corrections yield better motion artifact removal than either function alone (Di Lorenzo et al., 2019; Gao et al., 2022). A low-pass filter with a cutoff frequency of 0.14 Hz was applied to remove high-frequency noise (*hmr_BandpassFilt*). Optical density data were then converted to HbO_2_ and HHb concentrations via the modified Beer-Lambert law (*hmr_OD2Conc*), with a partial pathlength factor of 6 (Boas et al., 2004; von Lühmann et al., 2020). Any remaining motion artifacts were addressed by applying the gait-specific correlation-based signal improvement (CBSI) method (*hmr_MotionCorrectCbsi*) (Cui et al., 2010). While the CBSI method is most effective for calculating an activation signal that combines HbO_2_ and HHb, it also applies a correction (albeit nominal) on both the HbO_2_ and HHb signals separately. Applying this additional motion artifact correction step ensured a more conservative motion artifact correction approach, as previously used (Martini et al., 2024; Vitorio et al., 2020). A short-separation channel regression was performed to remove superficial hemodynamic responses from the four channels of interest (bilateral PPC). A baseline correction was performed: the mean of the signal from the middle 10 seconds of the 20-second quiet period was subtracted from the 3-minute walking period. We excluded the first 5s of the baseline period to avoid any residual activity following the earlier verbal instruction. We excluded the last 5s (i.e., immediately prior to gait initiation) to avoid anticipatory responses, as others have done (de Belli et al., 2021). Finally, the four channels were median averaged, such that the outcome measures comprised the median relative changes in HbO_2_ (ΔHbO_2_) and HHb (ΔHHb) concentration during each gait condition. The median relative ΔHbO_2_ and ΔHHb concentration was preferred to the mean, as it would be less sensitive to a potential long tail of the hemodynamic response over the 3-minute walk.

### Statistical analysis

Repeated measures ANOVAs were used to compare differences in the median relative ΔHbO_2_ and ΔHHb across gait conditions. Partial eta squared (*η*^*2*^) effect size was calculated and categorized: small (≥0.01), medium (≥0.06), large (≥0.14) (Cohen, 1988). Where *post-hoc* pairwise comparisons were made, Cohen’s *d* effect sizes were calculated and categorized: small (≥0.2), medium (≥0.5), large (≥0.8) (Cohen, 1988). Pearson’s correlation was used to determine the relationships between PPC ΔHbO_2_ and gait variability measures within each condition. Alpha was set a priori at <0.05. All statistical analyses were completed using JASP software (JASP 0.19.1; Amsterdam, Netherlands).

## Results

### Gait performance (Mean (SD)) in each condition

#### PPC activity

Results of a repeated measures ANOVA showed no significant effect of gait condition on PPC relative ΔHbO_2_, *F*_(2,40)_ = 2.36, *p* = 0.12. However, a medium effect size was observed (*η*^*2*^ = 0.11), driven by a more positive ΔHbO_2_ in the PPC during VC (Cohen’s *d* = 0.39) and VCP (Cohen’s *d* = 0.34) gait as compared to NC gait. PPC relative ΔHbO_2_ data, for each gait condition, are presented in Figure 3. Relative ΔHHb concentrations were not statistically different across gait conditions (NC, +0.04 ± 0.14μM; VC, -0.03 ± 0.15μM; VCP, -0.01 ± 0.14μM). Group averaged relative ΔHbO_2_ and ΔHHb across the 3-minute walking period, for each condition, are presented in Supplemental Figure 1.

**Fig 3:**
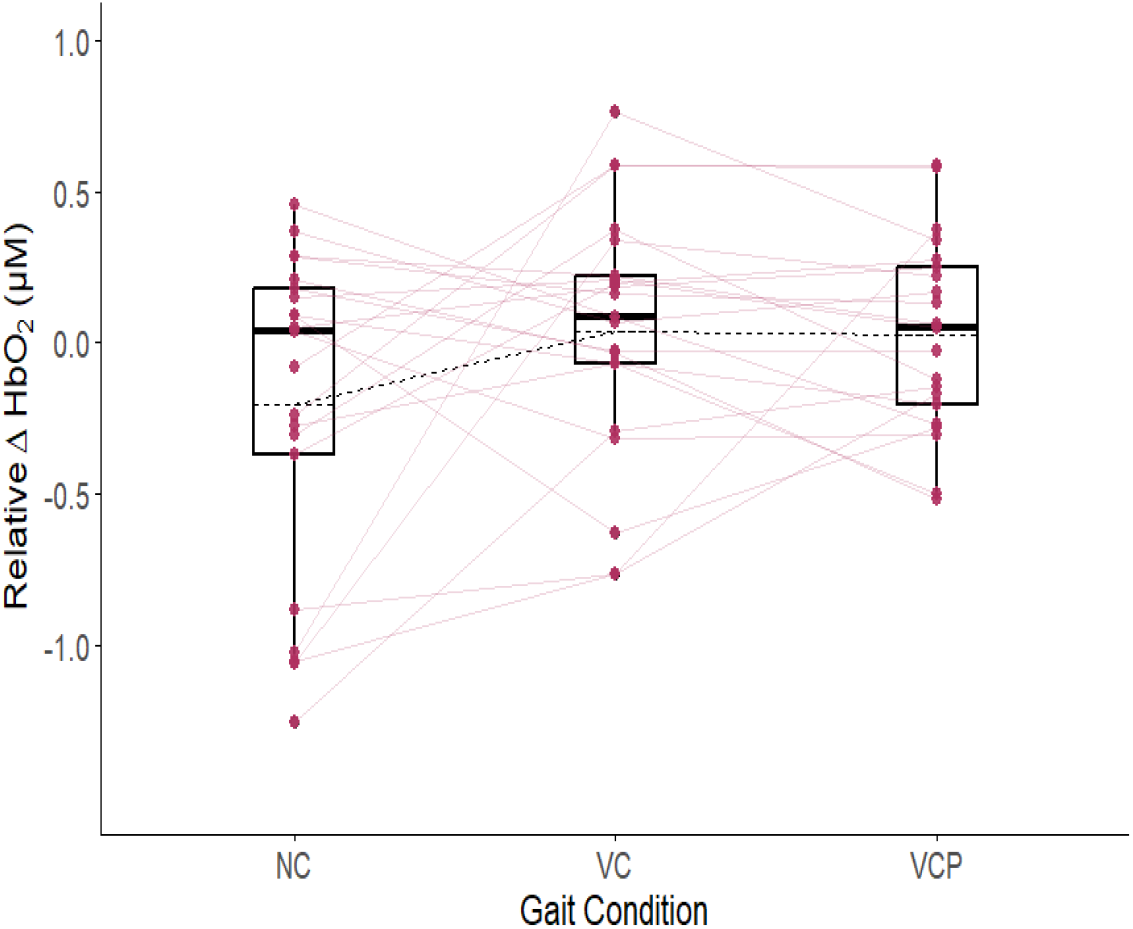
Box and scatter plots of the relative change in oxygenated hemoglobin (Δ HbO_2_), from baseline quiet standing, during each gait condition: NC, non-cued; VC, visually cued; VCP, visually cued with perturbations. Solid lines represent the median and dashed lines represent the mean for each condition

#### Associations between PPC activity and gait variability

Gait performance during each condition is presented in Table 1. Stride time variability significantly related to the relative ΔHbO_2_ in the PPC during the VC condition (Pearson’s *r* = 0.58, *p* = 0.01), such that higher stride time variability related to increased PPC activity (Figure 4). During VCP gait, stride time variability was not significantly related to the relative ΔHbO_2_ in the PPC. Neither stride length variability nor lumbar mediolateral RoM variability significantly correlated with the relative ΔHbO_2_ in the PPC during any gait condition.

**Table 1.** Gait performance measures during non-cued gait (NC), visually cued gait (VC), and visually cued gait with perturbations (VCP). * indicates a significant difference from NC gait. For VCP gait, † indicates a significant difference from VC gait.

| *n* = 21 | Gait Condition |  |  |
| --- | --- | --- | --- |
|  | NC | VC | VCP |
| **Gait speed (m/s) = 1.18 (0.16)** |  |  |  |
| Stride Time Variability (CoV %) | 2.04 (0.83) | 2.86 (0.53)* | 5.74 (1.03)*† |
| Stride Length Variability (CoV %) | 2.48 (0.92) | 3.01 (0.68)* | 7.63 (2.72)*† |
| Lumbar mediolateral RoM Variability (CoV %) | 13.01 (4.45) | 16.14 (6.97)* | 22.20 (10.77)*† |
| Stride Time (s) | 1.12 (0.07) | 1.06 (0.08)* | 1.06 (0.08)* |
| Stride Length (m) | 1.18 (0.13) | 1.12 (0.12)* | 1.12 (0.12)* |
| Lumbar mediolateral RoM (°) | 5.38 (1.68) | 4.69 (1.49)* | 5.12 (1.31)† |

**Fig 4:**
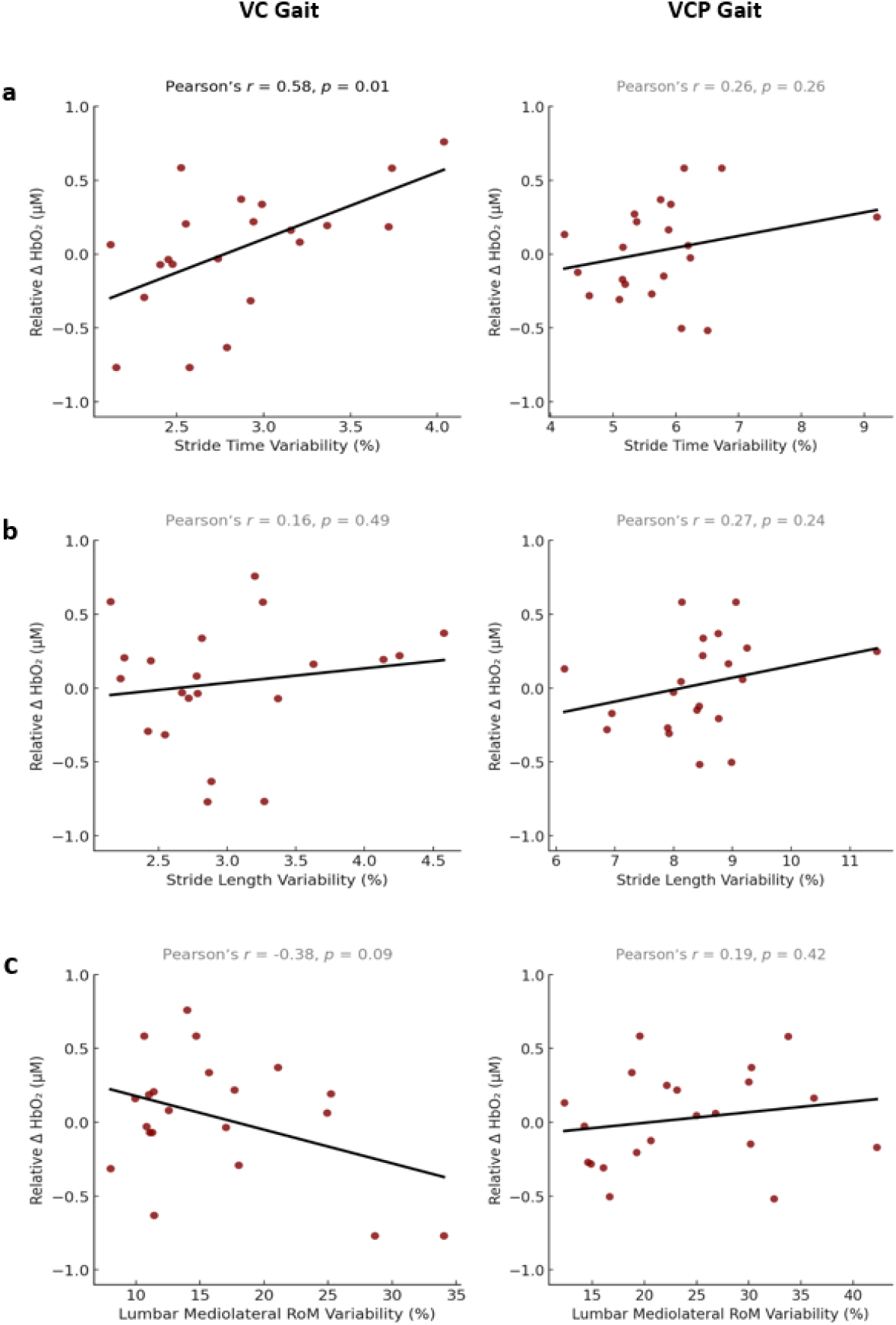
Scatter plots of the relationships between the relative change in oxygenated hemoglobin (ΔHbO_2_) in the PPC and the coefficient of variation for a: stride time, b: stride length, and c: lumbar mediolateral range of motion (RoM) during visually cued (VC) gait (left column) and visually cued gait with perturbations (VCP) (right column). Black lines represent the best fit lines

## Discussion

This preliminary study aimed to decipher PPC activity patterns during non-cued (NC) gait, visually cued (VC) gait and visually cued gait with perturbations (VCP). Our hypothesis for increasing PPC activity from NC to VC gait, and from VC to VCP gait, was not statistically supported, though we did observe moderate effect sizes to inform future study design. We also examined how PPC activity relates to gait variability during these gait conditions. Our hypotheses regarding relationships between PPC activity and gait variability were partially supported. Specifically, increased PPC activity (more positive ΔHbO_2_) significantly related to increased stride time variability, though during VC gait only. These results add to our understanding of cortical activity during gait and serve as a foundation for future studies involving populations with mobility deficits.

Moderate effect sizes were observed for PPC activity increases from NC to both VC and VCP gait among this sample of young adults. While not statistically significant, this increase in PPC activity with visual task complexity aligns with results from other studies that recorded PPC activity during similar VC gait tasks (Liu et al., 2024; Yokoyama et al., 2021). EEG studies report reduced alpha power in the PPC during VC gait compared to NC gait (Wagner et al., 2014; Yokoyama et al., 2021). Alpha power negatively correlates with blood-oxygen level-dependent-signal changes, which suggests that reduced alpha power indicates greater cortical activity (Moosmann et al., 2003). Further, Liu et al. (2024) demonstrated that during treadmill gait, PPC alpha power decreases as terrain unevenness increases. Though increasing terrain unevenness increases reliance on visual processing for guiding step placements (Liu et al., 2024; Matthis et al., 2018), somatosensory feedback from the uneven terrains may be a factor in the observed PPC activity. Across these VC gait paradigms, young adults leveraged swift and accurate visual processing, which is critical in complex gait environments. Unfortunately, visual processing slows with age (Ebaid & Crewther, 2019), underscoring the importance for studies quantifying PPC activity during VC gait in older adults. Results from these studies would establish the effect of slowed visual processing on gait impairment and fall risk.

While PPC activity appears to increase from NC to both VC and VCP gait, PPC activity levels during VC and VCP gait were not greatly elevated above those of baseline quiet standing. The non-significant increase was driven by a negative ΔHbO_2_ in the PPC during NC gait, i.e., PPC deactivation during walking relative to the brief standing baseline period (Figure 3). This pattern of PPC deactivation indicates that among young adults, PPC engagement was not necessary for NC treadmill gait, potentially because treadmill gait does not require significant visuomotor integration due to limited visual flow. Consistent with our findings, Lau et al. (2014) reported reduced cortical sensorimotor network involvement in young adults during treadmill walking compared to standing. Young adults typically exhibit gait automaticity (Clark, 2015), relying more on subcortical neural networks for generating the appropriate muscle activation patterns during gait. By contrast, standing requires considerable active cortical control for maintaining balance and posture (Vuillerme & Nafati, 2007). Combined, our results and previous evidence suggest that the PPC deactivation observed during NC gait relates to a shift toward subcortical gait control in young adults.

We did not observe a condition effect for PPC activity when comparing VC and VCP gait (Cohen’s *d* = 0.04). Other supraspinal structures involved in locomotor control (e.g., brainstem and cerebellum) may be more critical for gait modifications in response to rapidly shifting stepping target positions (Hoogkamer et al., 2017). Alternatively, processing in higher-order cortical regions beyond the PPC (e.g., PFC) could be more important during VC and VCP gait. Supporting this notion, Corporaal et al. (2018) reported that greater stepping accuracy during VC gait was associated with increased white matter tracts connecting attentional cortical regions (e.g., parietal and prefrontal cortices). Importantly, comparing NC to VC gait with targets presented at either fixed or variable step lengths, Le et al. (2023) reported no difference in PPC activity between conditions. However, these authors identified increased functional connectivity between the PPC and PFC during VC gait with variable step lengths. PPC-PFC connectivity serves as a key pathway of the frontoparietal network, which promotes externally oriented attention relevant for visuomotor performance when responding to unexpected stimuli (Menon & D’Esposito, 2022). Specifically, VC gait requires allocation of attention to external information (i.e., the positions of visual cues), highlighting a key role for the frontoparietal network. However, the absence of a significant increase in PPC activity from VC to VCP gait in our study, combined with the increased PFC activity observed during obstacle negotiation (Mirelman et al., 2017) and VC gait (Koenraadt et al., 2014), suggests that the PFC may comprise the dominant node of the frontoparietal network for VC gait performance in young adults.

The positive relationship between PPC activity and stride time variability observed during VC gait could be interpreted from two, somewhat opposing perspectives. Increased PPC activity from NC to VC gait may reflect the deployment of more neural resources in response to the increased environmental visuospatial processing demands of the task. This top-down strategy may be detrimental to gait rhythmicity. From this perspective, higher stride time variability reflects an unstable gait pattern, as observed in older adult and clinical populations (Hausdorff et al., 2001; Lord et al., 2011). This interpretation aligns with the Compensation-Related Utilization of Neural Circuits Hypothesis (Reuter-Lorenz & Cappell, 2008), and mirrors that offered for the positive association between increased cortical activity and gait variability among older adults (Nóbrega-Sousa et al., 2020). Concurrently quantifying PFC and PPC activity during our gait paradigm could provide evidence to support this potential explanation. Alternatively, increased PPC activity could be an adaptive mechanism allowing for flexible step timing adjustments for young adults, mirroring results from other species (Marigold et al., 2011). From this perspective, higher stride time variability during VC gait reflects a skillful adaptation strategy (Stergiou & Decker, 2011), where ongoing modulation of step timing supports stepping precision (Koenraadt et al., 2014). Higher PPC activity in response to the visuospatial processing demands may facilitate this strategy. Quantifying step accuracy will help to clarify the effect of higher PPC activity on gait performance during VC gait.

Notably, although stride time variability increased from VC to VCP gait, a positive relationship between PPC activity and stride time variability did not persist in the VCP condition. That PPC activity was similar during both VC and VCP gait may suggest that for these young adults, the visuospatial processing demands of both conditions were comparable. VCP gait imposes a degree of gait variability, as frequent step adjustments are necessary for good task performance. Applying the above argument for increased PPC activity as an adaptive mechanism supporting step accuracy, it would be reasonable to expect PPC activity to remain positively correlated with stride time variability during VCP gait. However, VCP gait performance may rely more on executive functions (e.g., attentional control, decision making) than VC gait. As previously suggested, the PFC potentially plays a pivotal role during VCP gait, and PFC activity in this condition conceivably relates to gait variability. Relatedly, Mirelman et al. (2017) reported a positive correlation between PFC activity and gait variability among older adults during obstacle negotiation. Assessing PFC activation during our gait paradigm will help to clarify the role of executive control in gait adaptability to shifting visual cues.

That we observed a relationship between PPC activity and temporal, but not spatial, gait variability aligns with the brain map of gait variability put forward by Tian et al. (2017). Structural MRI findings suggest that PPC grey matter volume negatively correlates with stride time variability in older adults (Beauchet et al., 2014). Using our paradigm to examine associations between real-time PPC activity and spatiotemporal gait variability in older adults will therefore be illuminating. Moreover, Pizzamiglio et al. (2018) indicated a role for the PPC in mediolateral gait control, specifically that higher PPC activity related to lower mediolateral center of mass motion during unperturbed gait. Our findings offer some support for this, as a moderate correlation (*r* = -0.38) emerged between higher PPC activity and lower lumbar mediolateral RoM variability during VC gait. Furthermore, lumbar mediolateral RoM was significantly lower during VC gait than NC gait. These findings indicate that in response to predictable visual cues during gait, young adults promote stability by restricting movement in the mediolateral direction, and this adaptive response may be supported by visuomotor integration in the PPC. Notably, evidence suggests that for older adults, the control of dynamic balance is more challenging in the frontal plane than in the sagittal plane (Vistamehr & Neptune, 2021). Accordingly, more falls occur in the frontal plane than in the sagittal plane (Parkkari et al., 1999). Therefore, unravelling the PPC’s role in mediolateral gait control during VC gait is a critical step toward understanding and mitigating fall risk in older adults.

The results and interpretation of this preliminary study need to be considered along with the following limitations. First, as Beauchet et al. (2009) insist, caution must be exercised when interpreting gait variability. Depending on the circumstances, both low and high variability of gait parameters may reflect good gait performance. As discussed, high variability of gait parameters may be considered a marker of adaptability to the gait environment. Future studies accounting for step accuracy could provide additional context to the present findings. Second, cortical areas beyond the PPC may be implicated in VC gait control, including the PFC (Le et al., 2023), premotor cortex (Wang et al., 2008), and supplementary motor area (Koenraadt et al., 2014). A more comprehensive assessment of activity across the cortex during our gait conditions could offer greater insights into the cortical mechanisms underlying VC gait performance. Additionally, the hemodynamic response delay following neural activity renders fNIRS unsuitable for assessing cortical activation changes within different phases of the gait cycle, and for a step- or stride-level comparison. Where reactive step adjustments are required, characterizing intra-stride neural dynamics with EEG would help to uncover how the precise timing of cortical contributions supports gait performance. Our VCP gait task design, involving target position shifts in different directions every 3-7 steps, meant it was not feasible to examine a potential effect of step adjustment direction on PPC activation. Finally, using the 10-20 system and visual inspection to identify channels corresponding to our cortical regions of interest is not the most rigorous approach, but has been successfully implemented across neuroimaging techniques (Herwig et al., 2003; Koenraadt et al., 2014; Shafiul Hasan et al., 2020; Velu & de Sa, 2013).

## Conclusion

This preliminary study offers new insights into PPC activity during VC treadmill gait in young adults. The moderate cue condition effect observed for PPC activity suggests that whether reactive step adjustments are necessary or not, greater PPC recruitment is required for VC gait than for NC gait. This likely reflects the increased visuomotor processing demands posed by visual cues. The positive relationship between PPC activity and stride time variability observed during VC gait potentially suggests that higher PPC activity supports modulation of step timing in response to visual stepping targets. Considering the association between high gait variability and fall risk among older adults (Hausdorff et al., 2001), future studies should extend these findings to aging populations. With evidence suggesting an important role for the frontoparietal network during VC gait (Corporaal et al., 2018; Le et al., 2023), expanding the cortical assessment to include the PFC should offer valuable insights. Quantifying PPC and PFC activity during VC and VCP gait in older adults, and relating cortical activity to gait performance, could help to uncover neurophysiological signatures of increased fall risk.

## Data & Code Availability

As this study is part of an ongoing, federally funded investigation, access to data and code is subject to restrictions. Access may be provided following a formal request.

