## Supplemental Figure 1 for "Posterior parietal cortex activity during visually cued gait: a preliminary study"

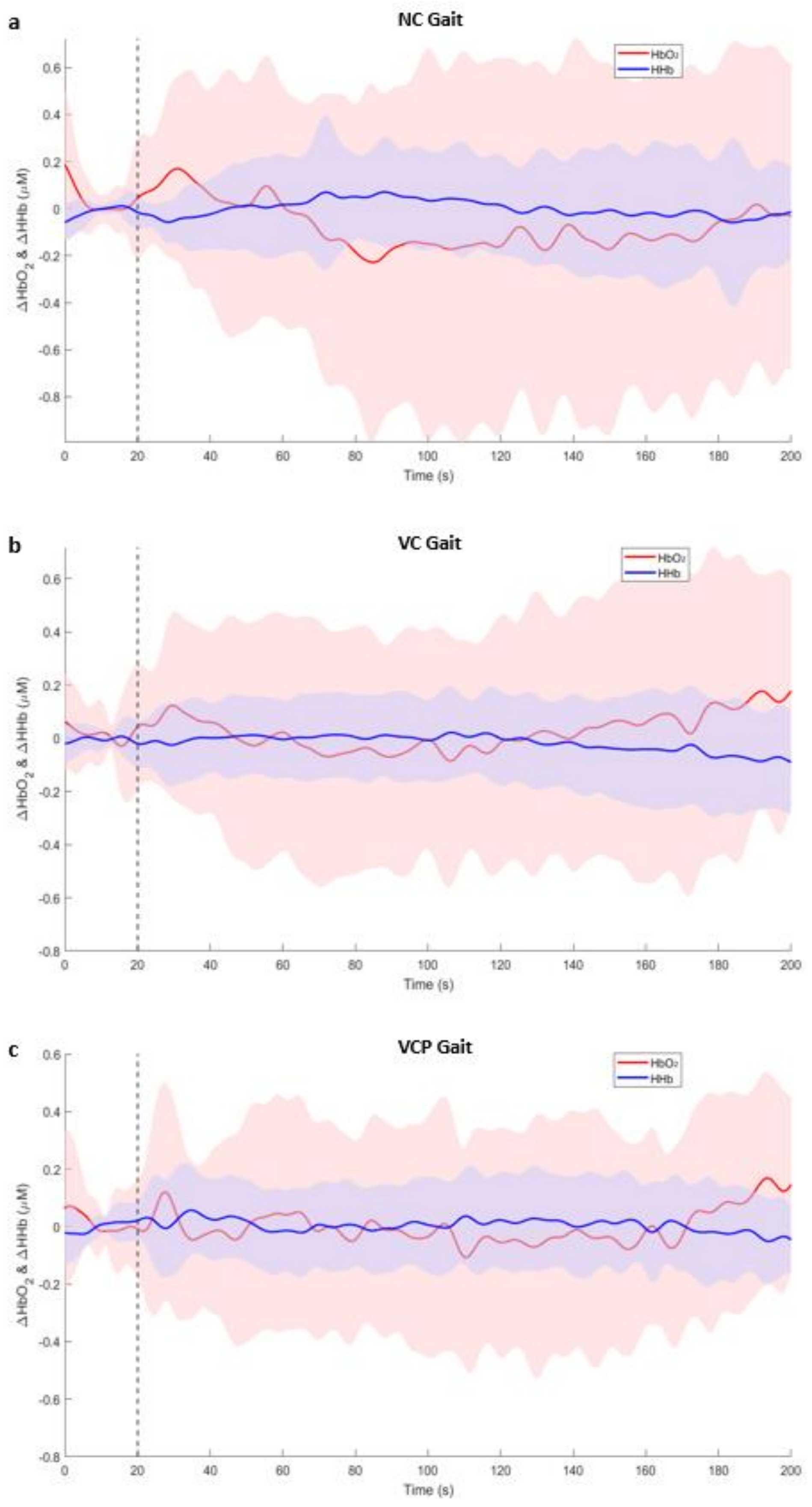
a
NC Gait
HbO2
HHb
ΔHbO2 & ΔHHb (μM)
Time (s)
b
VC Gait
HbO2
HHb
ΔHbO2 & ΔHHb (μM)
Time (s)
c
VCP Gait
HbO2
HHb
ΔHbO2 & ΔHHb (μM)
Time (s)

**Supplemental Fig.1** Group averaged relative $\Delta HbO_2$ and $\Delta HHb$ across the 3-minute walking period, for a: non-cued gait (NC), b: visually cued gait (VC); and c: visually cued gait with perturbations (VCP). Vertical dashed lines represent the end of the baseline standing period and the beginning of the walking period
